# A novel imaging biosensor for the detection of reversed replication forks and four-way junctions in human cells

**DOI:** 10.64898/2026.08.23.746510

**Authors:** Eva Malacaria, Elena Roscioli, Annapaola Franchitto, Pietro Pichierri

**Author notes:** Author to whom correspondence should be addressed: Pietro Pichierri.

## Abstract

Replication fork reversal and recombination-dependent replication protect perturbed forks by forming four-way DNA junctions. Despite their central role, detecting these intermediates in intact cells has historically required electron microscopy of bulk extracted DNA. Here, we describe a genetically encoded biosensor for four-way junctions based on nuclear-targeted bacterial RuvA tagged with GFP or Spot-tag. Combined with SIRF, RuvA selectively accumulates at nascent DNA following hydroxyurea- or camptothecin-induced replication stress. Recruitment requires fork-reversal factors and is abolished by a non-binding mutant (K84E/K119E), confirming specificity. The biosensor dynamically tracks reversed fork abundance, capturing MRE11-mediated fork degradation in BRCA2- or RAD52-deficient cells and its functional rescue. Finally, RuvA reveals RAD51-dependent four-way junctions at nucleolar rDNA arrays in unperturbed and stressed cells. This tool converts a bulk population measurement into a visualizable readout with single-cell and subnuclear resolution, enabling direct spatial analysis of DNA structures in situ.

## INTRODUCTION

The response to replication stress (RS) is of paramount importance for the maintenance of genome integrity and it comprises different steps, from sensing of the perturbed replication forks to spreading of the signal in the cell for cell cycle regulation or cell death engagement if excessive loads of DNA damage are generated at blocked forks ^1–8^. One of the steps of this multifaceted response is the remodelling of DNA at stalled forks with the formation of a reversed fork (RF) through a mechanism called fork reversal ^9–12^. Fork reversal catalyses the conversion of the Y-shaped DNA structure at replication forks into an X-shaped four-way DNA structure resembling a Holliday junction (HJ) in shape ^9,12^. This reaction involves the reannealing of the two nascent strands extruded from the templates through the action of a branch-migrating DNA annealing activity, which has its prototype in the bacterial RecG enzyme ^13,14^. Formation of a RF serves two functions: masking vulnerable ssDNA at the fork and moving any DNA damage residing in the ssDNA template back into a dsDNA configuration for repair. Last, RFs additionally provide the cells with a versatile intermediate that – by resembling a HJ – can be used for recombination-based pathways of fork restart ^9,12^. In human cells, RF formation can be performed by several proteins, which are key for fork recovery and genome integrity, including SMARCAL1 – the central RF-relevant enzyme – ZRANB3, HLTF and also the recombinase RAD51 ^9,15–20^. Other proteins can stimulate fork reversal, suggesting that either cells developed a striking redundancy or there is specificity in terms of DNA environment ^9,10^. Fork reversal is a physiological event, critical whenever cells experience replication fork perturbations, but is also a phenomenon that needs to be regulated. Indeed, deregulated engagement of fork reversal with a pathological upregulation can lead to DNA damage or fork degradation as it has been observed in the absence of proper control of SMARCAL1 or of antagonists such as RADX and RAD52 ^21–24^.

Although the number of RFs in unperturbed cells is very low – estimated in about the 5% in transformed cells – it clearly increases under RS approaching the 30% as it can be inferred from electron microscopy (EM) ^11,25^. This figure is probably underestimated, since RFs must survive psoralen crosslinking and DNA extraction to be scored, and since RFs are dynamic structures that can be restored or degraded before the sample is fixed. Moreover, not all the agents inducing RS are equally good in stimulating RFs. For instance, low-doses of the Topo-I poison camptothecin (CPT) are strong inducers of RFs ^26,27^.

Reversed forks are also cancer-relevant structures since their stability relies on the presence of recombination enzymes such as RAD51 and the two cancer-predisposition genes BRCA1 and BRCA2 ^28–33^. Deprotection of RFs leads to degradation of the two extruded and reannealed nascent strands by exonucleases activity of MRE11 and EXO1 and DSBs formation by SLX4/MUS81 ^34^. Of note, degradation of RFs ditches the four-way junction converting it into a DSB and contributes to PARPi and chemosensitivity of BRCA-deficient cancer cells ^35–38^. This implies that estimating the number of RFs and/or their propensity to being degraded may be a biomarker for therapeutic purposes.

Currently, the presence of RFs can be quantified directly only using purification of DNA intermediates and visualization by EM ^39^. EM remains the only method that directly visualizes reversed forks, but it requires a large number of cells, extreme processing of the sample and is an ensemble technique. Moreover, EM provides no spatial information, no genomic context within the nucleus and no single-cell resolution. Tools that can directly visualize the presence of RF at the single-cell level are not available, while only surrogate approaches, such as presence of SMARCAL1 at the fork, the unveiling of nascent ssDNA at the fork or fork rates estimated by DNA fibers, can be used for this purpose. However, all these approaches suffer from potential confounders: the presence of SMARCAL1 does not tell if the protein is active or performed its job, nascent ssDNA is also an intermediate of DSB processing, and fork rate can be affected by several factors and is completely blind to dynamics and spatial information. Therefore, questions that are routine in cell biology remain unanswerable for this intermediate: whether RFs are distributed uniformly in the nucleus or concentrate in specific compartments, whether their load differs between individual cells of the same population, and which factors reside in their proximity.

In the literature, biosensors for other RS-relevant structures have been reported though. Catalytically-dead and imaging-enabled RNaseH1 is used to detect R-loops ^40^, providing a guide for a biosensor of RFs and four-way junctions at large. In bacterial cells, HJs are cleaved by the RuvABC complex ^41–43^. In this complex, RuvA represents the structure-binding module that embraces the four-way junction as a tetramer, in a sequence-independent manner, acting as a platform for the other two – enzymatically-active – proteins ^41–43^. No RuvBC is present in human cells, so expression of the RuvA alone would represent a potential biosensor of four-way junctions, including RFs, at the single cell level by imaging. Consistent with this, we previously reported that RuvA expressed in the nucleus of human cells is found in chromatin fractions, that its chromatin association increases upon treatment with CPT, and that it can shield four-way junctions from being cleaved by the GEN1 endonuclease ^44^. Of note, in the same study, ectopic RuvA expression was sufficient to increase micronuclei in unperturbed cells, indicating that stable engagement of junctions is at the same time the basis of detection and a potential perturbation of the very structures being reported – an aspect that we characterize directly here and that also provides a deliberate handle to interrogate the persistence of four-way junctions.

Here, we show that a codon-optimized, nuclear-targeted, RuvA with either a GFP tag or an independent tag – Spot-tag – can be found at perturbed replication forks by SIRF after HU and nanomolar doses of CPT. Fork recruitment is time-dependent, being higher when EM reported the higher number of RFs, is genetically dependent on the presence of SMARCAL1, ZRANB3 or RAD51, and is lost upon introduction of the K84E+K119E mutations known to abrogate HJ-binding, which also abolish accumulation of RuvA in chromatin. The presence of the RuvA biosensor at perturbed forks is reduced in a MRE11-dependent manner when BRCA2 is depleted and is upregulated in RAD52-deficient cells when MRE11 is inhibited. Of note, although no RuvA foci can be seen by confocal microscopy after treatment – suggesting that occupancy at each junction is stoichiometrically low – a clear nucleolar staining, overlapping with nucleolin, enhanced by HU and reduced by RAD51 inhibition, can be appreciated even in unperturbed cells, pointing to a constitutive formation of four-way junctions at the rDNA. In conclusion, our study reports the generation of a four-way junctions – including RFs – biosensor that can reveal the presence of those structures at the single cell level, allowing to determine where they are formed, when they interact with other factors and which factors, and to perform topological studies using super-resolution techniques such as dSTORM or expansion microscopy. Moreover, being an avid binder, the RuvA biosensor can also help in investigating the consequences of the persistence RFs, and of four-way junctions at large, in the cell, for which we provide proof of principle.

## METHODS

### Cell lines and culture conditions

The U2OS cells were obtained from ATCC and maintained in Dulbecco’s modified Eagle’s medium (DMEM; Life Technologies) supplemented with 10% foetal bovine serum (Boehringer Mannheim) and incubated at 37°C in a humidified 5% CO2 atmosphere. To generate inducible U2OS RuvA-SpotTag, cells were transduced with lentivirus expressing RuvA-SpotTag (Vectorbuilder: pLV[Exp]-Bsd-TRE3G> {NLS-ruvA GFP-spot-tag}) at increasing multiplicity of infection (MOI) and (VectorBuilder: pLV[Exp]-Neo-CMV>Tet3G) at 1 of MOI in a DMEM supplemented with 10% tetracycline free bovine serum (Aurogene). After puromycin selection at 1µg/ml, cells were used to perform experiments.

### Chemicals

Doxycycline was dissolved in DMSO at 1mg/mg and was added to the culture medium at a concentration of 1µg/ml for 48hrs prior to perform experiments. HU was added to a culture medium at 2mM from stock solutions 200 mM prepared in water to induce DNA replication arrest or DNA damage. The B02 compound (Selleck), an inhibitor of RAD51 activity, was used at 27 µM. CldU (Sigma-Aldrich) was dissolved in sterile water as a 200 mM stock solution and used at 50 μM. IdU (Sigma-Aldrich) was dissolved in sterile DMEM as a stock solution 2.5 mM and stored at −20°C. The RAD52 inhibitor, EGC (Sigma-Aldrich) was dissolved in DMSO at 100mM, and stock solution was stored at −80° and was used at 50µM. MIRIN, the inhibitor of MRE11 exonuclease activity (Calbiochem), was used at 50 µM. Camptothecin (ENZO Life Sciences) was dissolved in a DMSO at 10mM, and stock solution (25µM) was stored at -20° and used at 25nM.

### Transfections

SMARCAL1 siRNA was transfected at a final concentration of 40nM using interferin (Polyplus) 48 h before performing experiments. BRCA2 siRNA was used at 50 nM using interferin and experiments were performed after 48h post transfection. The NLS-RuvA-EGFP vector was transfected in MRC5 or U2OS cell lines using Neon transfection system (Life technologies) 48 h prior to perform experiments.

### Real-time PCR

To evaluate ZRANB3 mRNA expression following transient transfection with siRNA oligos, total RNA was extracted from U2OS cells and reverse-transcribed into cDNA (iScript cDNA Synthesis Kit (Bio-Rad). Quantitative real-time PCR was performed on an Applied Biosystems Real-Time PCR System (7500 series) using target-specific fluorogenic probes for ZRANB3 (TaqMan Hs00999251_m1). Human ACTB (Actin) was used as the endogenous housekeeping control for normalization. Relative mRNA expression levels were quantified using the comparative 2^-ΔΔCt^ method, and all reactions were performed in biological triplicates.

### Western blot analysis

Western blots were performed using standard methods. Blots were incubated with primary antibodies against anti-SpotTag (Chromoteck 1:1000), Lamin B1 (Abcam, 1:10,000), anti-GAPDH (Millipore, 1:5000, anti-SMARCAL1 (Bethyl 1:1000), anti-BRCA2 (Bethyl 1:1000) anti-GFP (Santa Cruz 1:1000) and anti-Vinculin (Santa Cruz 1:500). After incubations with horse radish peroxidase-linked secondary antibodies (Jackson Immuno Research, 1:30,000), the blots were developed using the chemiluminescence detection kit ECL-Plus (Amersham) according to the manufacturer’s instructions. Quantification was performed on blot acquired by ChemiDoc XRS+ (Bio-Rad) using Image Lab software, and values shown on the graphs represent a normalisation of the protein content evaluated through Lamin B1 or Vinculin immunoblotting.

### Single-cell assay for in situ protein interaction with nascent DNA (SIRF)

Exponential growing cells were seeded onto microscope chamber slide. On the day of experiment, cells were incubated with 125 µM EdU for 8 min and treated as indicated. After treatment, cells were pre-extracted in CSK buffer at 300 nM NaCl and fixed with 3% PFA, 2% sucrose in PBS 1X for 15 min at RT. Cells were then permeabilized with 0.25% TritonX-100 for 15 min and blocked in 3% BSA/PBS for 20 min. For the EdU detection was applied the Click-iT™ EdU Alexa Fluor™ Imaging Kit (Invitrogen) using 5mM Biotin-Azide for 30 minutes at RT. The primary antibodies used were as follows: mouse monoclonal anti-SpotTag (Chromoteck 1:2000), rabbit anti-biotin (Abcam, 1: 3000). The negative controls were obtained by using only one primary antibody or both in not induced cells. Samples were incubated with secondary antibodies conjugated with PLA probes MINUS and PLUS: the PLA probe anti-mouse PLUS and anti-rabbit MINUS (Navinci Diagnostics). The incubation with all antibodies was accomplished in a humidified chamber for 1 h at 37 °C. Next, the PLA probes MINUS and PLUS were ligated using two connecting oligonucleotides to produce a template for rolling-cycle amplification. After amplification, the products were hybridized with red fluorescence-labelled oligonucleotide. Samples were mounted in Prolong Gold antifade reagent with DAPI (blue). Images were acquired randomly using the Eclipse 80i Nikon Fluorescence Microscope, equipped with a Virtual Confocal (ViCo) system. The analysis was carried out by counting the SIRF spot for each nucleus. For quantitative imaging, chamberslides were imaged using an Evident SpinX HCS system using the ScanR software.

### Immunofluorescence

Cells were cultured onto 8-well chamber-slides (Millicell Sigma-Aldrich) and RuvA-Spot-Tag expression was induced or not with 1 µg/ml Doxycycline 48 h before fixation. Cells were treated for 4 hours with 2 mM Hydroxyurea. After treatment, cells were pre-extracted in 0.5% Triton X-100 for 10 min on ice, fixed with 3% PFA, 2% sucrose in PBS 1X for 15 min at room temperature (RT) and blocked in 5% bovine serum albumin (BSA) in PBS solution for 1 h at 37°C in a humidifier chamber. Staining with mouse monoclonal anti-Spot-Tag (1:500, ChromoTek 28A5) diluted in a 1%BSA/0,1% saponin in PBS solution, was carried out for 1h at RT in a humidifier chamber. After extensive washing with PBS, a species-specific fluorescein-conjugated secondary antibody (Alexa Fluor 488-conjugated Goat Anti-Mouse IgG (H + L), highly cross-adsorbed; Life Technologies) was applied for 1 h at 37°C in a humidifier chamber. Counterstaining was performed with 0.5 μg/ml 4,6-diamidino-2-phenylindole (DAPI). Secondary antibodies were used at 1:200 dilution. Slides were analysed with an Ecliplse 80i Nikon Fluorescence Microscope, equipped with a Video Confocal (ViCo) system at 40× magnification.

### Co-Immunofluorescence (CoIF) and confocal microscopy

Cells were cultured onto 8-well chamber-slides (Millicell Sigma-Aldrich) and RuvA-Spot-Tag expression was induced or not with 1 µg/ml Doxycycline 48 h before fixation. Cells were treated for 4 hours with 2 mM Hydroxyurea or for 1h with 25 nM Camptothecin in combination or not with a pretreatment with 27 µM RAD51 inhibitor B02 (Selleckchem, S8434) 30 min before treatment. After treatment, cells were preextracted in 0.5% Triton X-100 for 10 min on ice, fixed with 3% PFA, 2% sucrose in PBS 1X for 15 min at room temperature (RT) and blocked in 5% bovine serum albumin (BSA) in PBS solution for 1 h at 37°C in a humidifier chamber. Staining with rabbit monoclonal anti-Nucleolin (1:200, ABclonal A20910) diluted in a 1%BSA/0,1% saponin in PBS solution, was carried out over night at 4°C in a humidifier chamber. The following day, after further washing with PBS, staining with mouse monoclonal anti-Spot-Tag (1:500, ChromoTek 28A5) diluted in a 1%BSA/0,1% saponin in PBS solution, was carried out for 1h at RT in a humidifier chamber. After extensive washing with PBS, species-specific fluorescein-conjugated secondary antibodies (Alexa Fluor 488-conjugated Goat Anti-Mouse IgG (H + L) and Alexa Fluor 594-conjugated Goat Anti-Rabbit IgG (H + L), highly cross-adsorbed; Life Technologies) was applied for 1 h at 37°C in a humidifier chamber. Counterstaining was performed with 0.5 μg/ml 4,6-diamidino-2-phenylindole (DAPI). Secondary antibodies were used at 1:200 dilution. Slides were analysed with an Eclipse 80i Nikon Fluorescence Microscope, equipped with a Video Confocal (ViCo) system at 40× magnification. For the acquisition of Z-stacks, chamberslides were imaged using the Evident SpinX HCS system and the Cellsens software.

### Statistical analysis

Experiments shown are representative of at least three independent biological replicates unless otherwise indicated in the figure legend. Significance was assessed using the built-in tools in Prism 10 (GraphPad Inc.) by Kruskal-Wallis’ test followed by post-hoc Dunn test for FDR for experiments with more than two samples, and by the two-tailed Student’s t-test to evaluate the means from normal distributions when analysing two samples. P < 0.05 was considered as significant. Statistical significance was always denoted as follow: ns = not significant; *P < 0.05; **P < 0.01; ***P < 0.01; ***P < 0.001. If not otherwise indicated in the figure legend, analysis was performed by Kruskal-Wallis’ test followed by post-hoc Dunn. Specific statistical analyses are reported in the relevant legend. No statistical methods or criteria were used to estimate sample size or to include/exclude samples.

## RESULTS

### RuvA expression and cell line generation

RuvA is a part of bacterial complex RuvABC which is responsible for recognition of four-way junction as Holliday junction in bacteria ^41,43^. We previously demonstrated that when a nuclear-localisation signal-supplemented RuvA (NLS-RuvA) is transfected in human cells can localise in chromatin fractions and functionally interfere with binding of the structure-specific, HJ, endonuclease GEN1 ^44^. This suggested that ectopic RuvA may be used as four-way biosensor in the cells to visualize sites where these structures are formed or accumulate in response to RS using imaging. To this end, we improved the original NLS-RuvA-GFP construct by introducing few codon optimisations, including a start codon ATG and also generated an independent construct able to deliver inducible expression of an NLS-RuvA-Spot-tag version of the protein through lentiviral delivery (Fig. 1A). To specifically localise RuvA at perturbed replication forks, we decided to use the SIRF assay in transiently-transfected U2OS cells treated with 2 mM HU for increasing time. As expected, transient transfection efficiently expressed RuvA in the cells (Fig. 1B). As shown in Figure 1C few SIRF spots representing RuvA localisation at replication forks were found in untreated cells whereas the exposure to HU substantially increased their number. Of note, SIRF controls using each single antibody failed to reveal discrete spots (Figure 1C – image panel). To perform a detailed characterisation of the RuvA biosensor, we generated stable cells lines using the other RuvA construct that replaces EGFP with a linker sequence and a Spot-tag on top of it (referred to RuvA-Spot thereafter). This configuration kept the tag at the C-terminus of RuvA, representing the region naturally protruding out of the DNA-binding module for the association with RuvBC (Fig. 1A) but reduced the size of the RuvA chimera. RuvA-Spot was placed under a Tet3G promoter for the conditional induction by doxycycline (Dox). We infected U2OS cells, which are widely used as model system in the field, and tested different MOI (multiplicity of infection). Thus, we selected cells infected with MOI 0.5 to keep the overall level of RuvA in the range of endogenous proteins and facilitate the detection by SIRF. Using these cells and tested different time of Dox induction (Fig. 1D). Although RuvA-Spot started to be detectable by WB as soon as 4 hrs of Dox, we analysed the binding between Spot-Tag and EdU-labelled nascent DNA at fork after 48 hours of induction with Dox. We next induced cells for 48hrs and performed SIRF assay in cells treated or not with 2 mM HU at 2 and 4 hrs and with 25 nM CPT for 1 hr. Consistent with the RuvA-SIRF from transient-transfected cells, spots were detected already at 2 hrs of treatment and increased at 4 hrs (Fig. 1E). Of note, and in agreement with EM data on RFs ^16^, the low dose of CPT was more efficient than HU in inducing RuvA recruitment (Fig. 1E). SIRF spots were barely detectable in both Dox-induced cells in which PLA was performed using only each single antibody and without Dox – that is in absence of RuvA (Fig. 1E).

**Figure 1.**
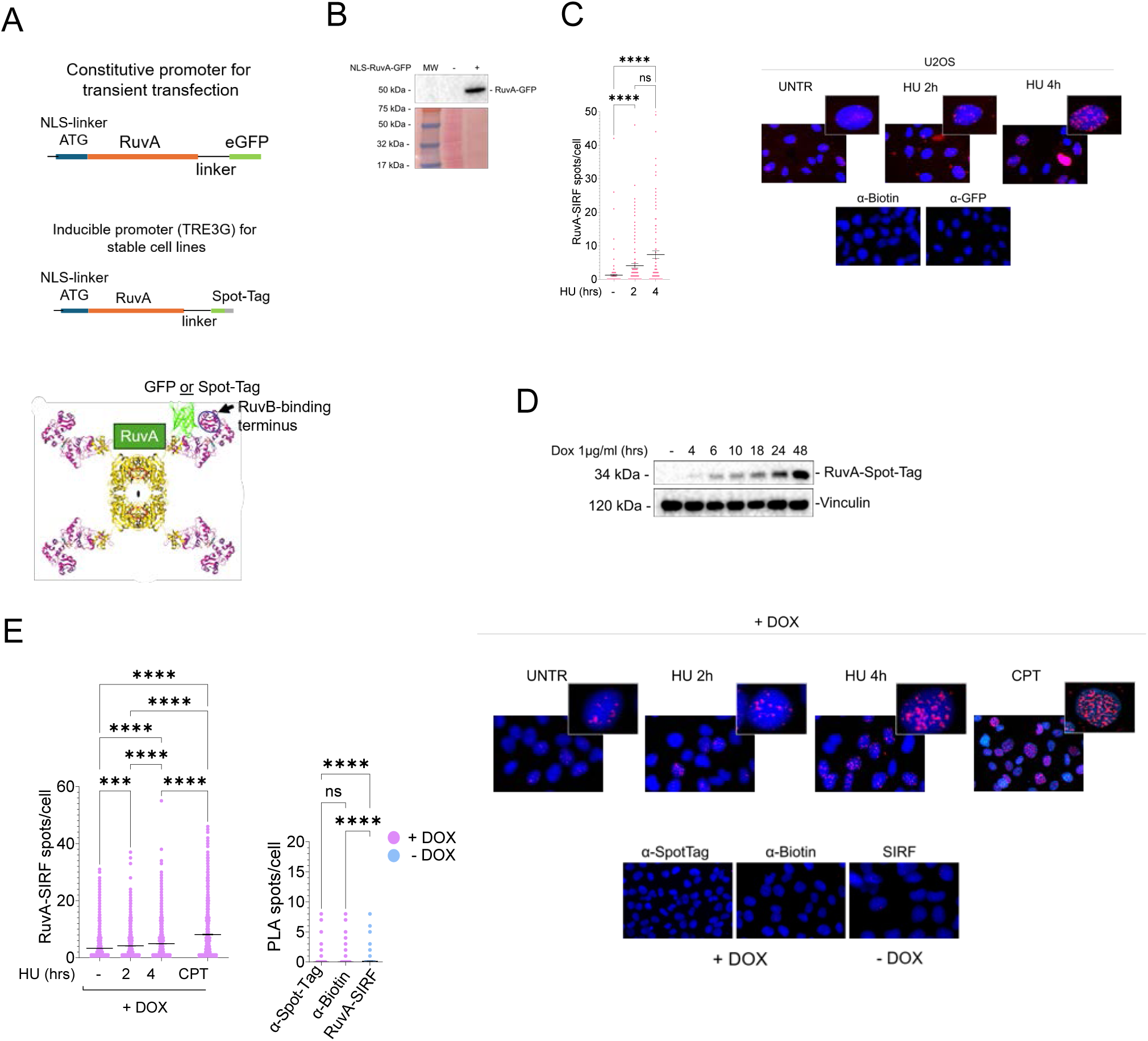
RuvA is recruited to stalled replication forks. **(A)** Cartoon representing the modular structure of the NLS-RuvA construct and the arrangement of the two biosensors. The scheme of RuvA shows that the tags are placed in the exposed region, where the protein engages with RuvB. **(B)** Western blot (top) shows RuvA-GFP expression levels; Ponceau S staining served as a loading control. **(C)** In situ protein interactions at nascent DNA (SIRF) assay in cells transfected with NLS-RuvA-GFP 48 h prior to analysis. SIRF was performed using anti-biotin and anti-GFP antibodies. Graph shows the number of spots per cell from combined experiments (N=3). Representative fluorescence images are shown (right). Data includes mean ± SEM; ns = not significant, \*\*\*\**PP* < 0.0001 by Kruskal–Wallis test. Representative control images showing single-antibody PLA controls (anti-biotin only for EdU, or anti-GFP only). **(D)** Western blot analysis of RuvA-Spot-Tag expression at indicated time points following induction with 1 µg/ml doxycycline (DOX) for 48 hrs. Vinculin served as a loading control. **(E)** SIRF assay evaluating RuvA recruitment after 48 hrs of DOX induction. Quantification was performed using an Olympus ScanR High-Content Imaging System (> 1000 cells analyzed per condition from N=3). Graph shows the number of spots and the mean PLA spots per cell ± SEM; \*\*\**PP* < 0.001, \*\*\*\**PP* < 0.0001 by Kruskal–Wallis test. Single-antibody controls (anti-biotin only and anti-Spot-Tag only) SIRF controls from non-induced cells, and representative images are shown.

Altogether our data indicated that RuvA-GFP and RuvA-Spot can localise at nascent DNA both in untreated and in HU-treated cells. Upon fork stalling, both RuvA biosensors increase their localisation at nascent DNA possibly representing four-way junctions *in situ*.

### Downregulation of fork reversal substantially prevents RuvA recruitment

Having shown that the RuvA biosensor can be localised at nascent DNA, we next wanted to define if this localisation actually marks RFs or four-way junctions at large. In response to RS, formation of these structures is genetically modifiable by removing the factors contributing to their formation. Several proteins are involved in fork reversal: SMARCAL1, RAD51, FBH1, ZRANB3, HLTF and their regulation is important to guarantee fork remodelling, while HJ largely depends on RAD51 ^9^. Thus, we downregulated some key fork reversal proteins as SMARCAL1 and RAD51 to assess if RuvA-Spot localisation at perturbed forks was affected. The expected result was that, if RuvA SIRF spots really marked RFs and/or HJ, then labelling should have been reduced in depleted cells. As shown in Figure 2A, RuvA-Spot increased its association with nascent DNA as detected by SIRF assay upon replication fork perturbation with 2 mM HU for 4 hrs (Fig. 2A). Of note, depletion of SMARCAL1 greatly suppressed SIRF spots as RAD51i did, and the extent of suppression was roughly similar (∼ 70 %). Combination of SMARCAL1 depletion and RAD51i increased suppression of the RuvA-Spot fork-associated signals (Fig. 2A). Consistent with this result, also depletion of ZRANB3 – another protein involved in RFs generation – resulted in a substantial reduction in the number of SIRF spots of RuvA-Spot (Fig. 2B). Biotin-Biotin control PLA demonstrated that RuvA-Spot expression does not affect EdU incorporation excluding that the observed, RS-dependent, increase in RuvA-Spot SIRF signal was unrelated to actual accumulation at perturbed forks (Fig. S1). The observed reduction in RuvA-Spot binding at nascent DNA after RS when nucleofilament assembly of RAD51 – the main human recombinase – is inhibited, supports the specific detection of four-way junctions represented by HJs and RFs, which requires RAD51 to be formed ^9,45^. To better appreciate the contribution of HJs, which are thought to largely derive after fork collapse, we performed a neutral Comet assay in cells expressing RuvA-Spot in the presence of HU at early time- points when, normally, they are barely formed in wild-type cells. As shown in Supplementary Figure 2, a very low-level of DSBs was detectable at the early time-point of HU or 25 nM CPT treatment used for SIRF assay irrespective of the expression of the RuvA-Spot, while DSBs were readily detected in the positive control represented by the 5 µM CPT.

**Figure 2.**
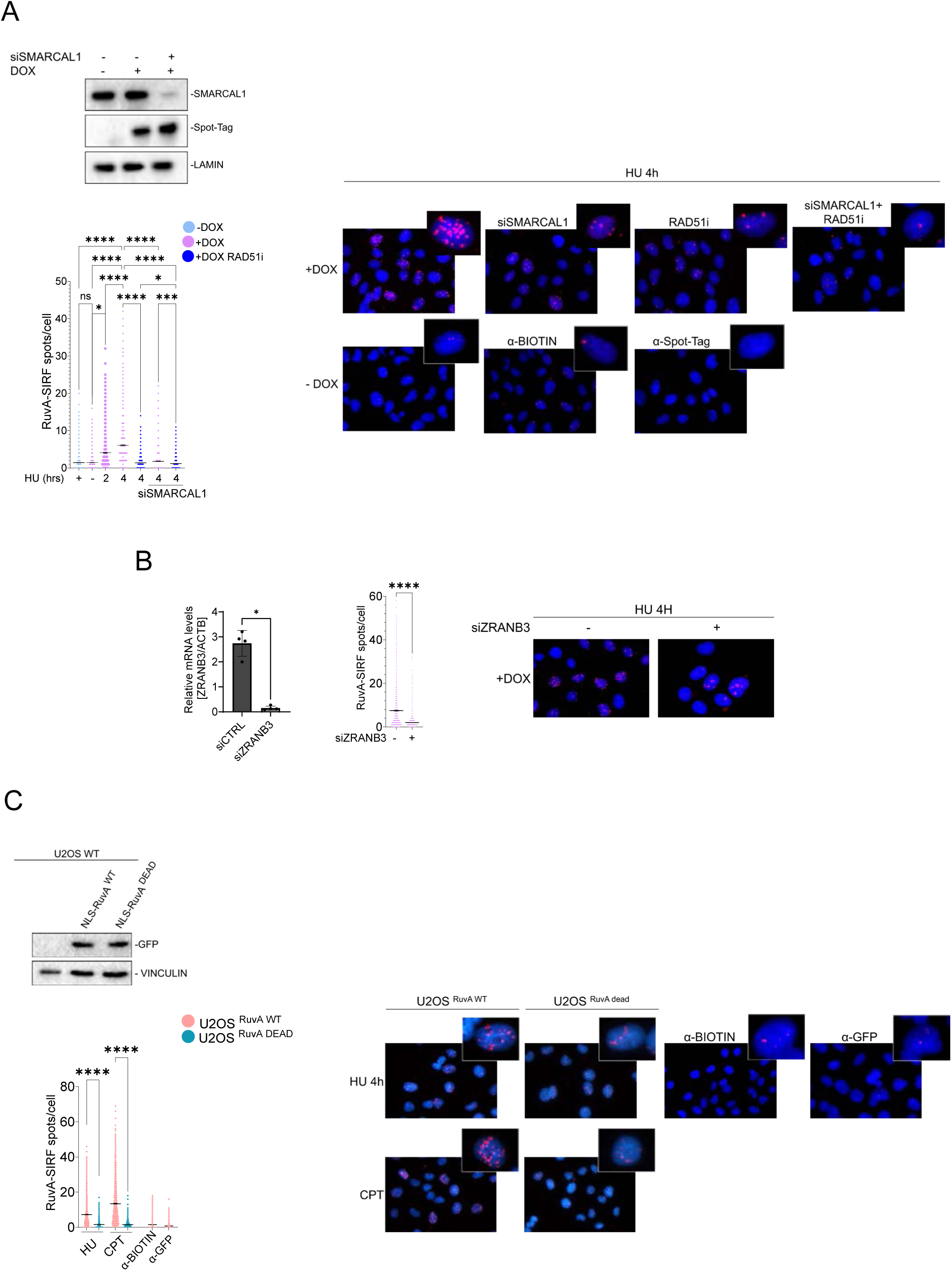
Downregulation of fork reversal factors impairs RuvA recruitment at reversed forks. **(A)** Cells were transfected with non-targeting control siRNA (siCTRL) or siRNA targeting *SMARCAL1*. Western blot analysis shows depletion efficiency; Lamin B1 served as a loading control. Quantification of PLA spots per cell was performed using an Olympus ScanR High-Content Imaging System (> 1000 cells per condition; N=3). Graph shows the number of spots and the mean PLA spots per cell ± SEM; ns = not significant, *PP* < 0.01, \*\*\*\**PP* < 0.0001 by Kruskal–Wallis test. **(B)** Analysis of RuvA recruitment in cells transfected with siCTRL or siRNA targeting *ZRANB3*. Real-time qPCR showed efficacy of depletion relative to *ACTB* control. In situ PLA analysis of DNA–RuvA interaction was conducted following indicated treatments. Quantification (> 1000 cells per condition; N=3) shows the number of spots and the mean PLA spots per cell ± SEM; \*\*\*\**PP* < 0.0001 by Kruskal–Wallis test. Representative images are shown. **(C)** Comparative recruitment of wild-type (NLS-RuvA-WT-GFP) and junction-binding-deficient (NLS-RuvA-DEAD-GFP) variants. Top panel: Western blot showing protein expression 48 hrs post-transfection. Graph shows the number of PLA spots per cell following indicated treatments (> 1000 cells per condition; N = 3). Negative controls incubated with single antibodies (anti-biotin or anti-GFP only) are included. Data includes mean ± SEM; \*\*\*\**PP* < 0.0001 by Kruskal–Wallis test.

We subsequently exploited specific mutations in RuvA shown to abrogate its binding affinity for four-way junctions ^46^. To investigate this, we performed K84E+K119E site-directed mutagenesis on the NLS-RuvA-GFP plasmid and utilized the resulting construct to transfect the U2OS cell line. To evaluate the capacity of mutated RuvA to interact with reversed forks, we employed the SIRF assay, quantifying RuvA recruitment in comparison to the wild-type protein upon RS. Our analysis demonstrated that K84E+K119E mutations, which disrupt the ability of RuvA to bind four-way junctions ^46^, lead to a large decrease in RuvA recruitment at nascent DNA as evidenced by SIRF assay (Fig. 2D). These data demonstrate that the RuvA detected by SIRF is specifically recognizing four-way-junctions.

Collectively, these results indicate that ectopically-expressed RuvA is detecting DNA structures formed after RS in a SMARCAL1, RAD51 or ZRANB3-dependent mechanism. Together with the suppression of the signal by the HJ binding-dead mutant RuvA, our findings demonstrate that RuvA, by a recognizing HJ-like junctions at the perturbed forks can be effectively utilized as a high-fidelity biosensor for detecting reversed forks and four-way junction at large *in vivo*.

### The RuvA biosensor can detect pathological transactions at reversed forks

Having shown that RuvA is a RF biosensor *in vivo*, we next wanted to evaluate its ability to detect changes in the abundance of RFs occurring when the stability of these structures is undermined by pathological conditions such as in BRCA2-deficient cells ^47^. In the absence of BRCA2, RFs are unstable, get degraded by MRE11 and EXO1, and eventually degenerate into MUS81-dependent DSBs ^34^. This is reflected by reduced number of RFs detected by EM, which can be rescued upon MRE11 inhibition ^31,32,34^. To evaluate how RuvA engagement on RFs was affected in this context, we executed a SIRF assay in the BRCA2-depleted RuvA-inducible cells after HU treatment and in the presence or absence of the MRE11 inhibitor, MIRIN. Consistent with EM data, depletion of BRCA2 was found to reduce the recruitment of RuvA to reversed replication forks detected by SIRF (Fig. 3A). This recruitment deficiency was restored through treatment with MIRIN (Fig. 3A), recapitulating what reported using EM ^29,31,34^.

**Figure 3.**
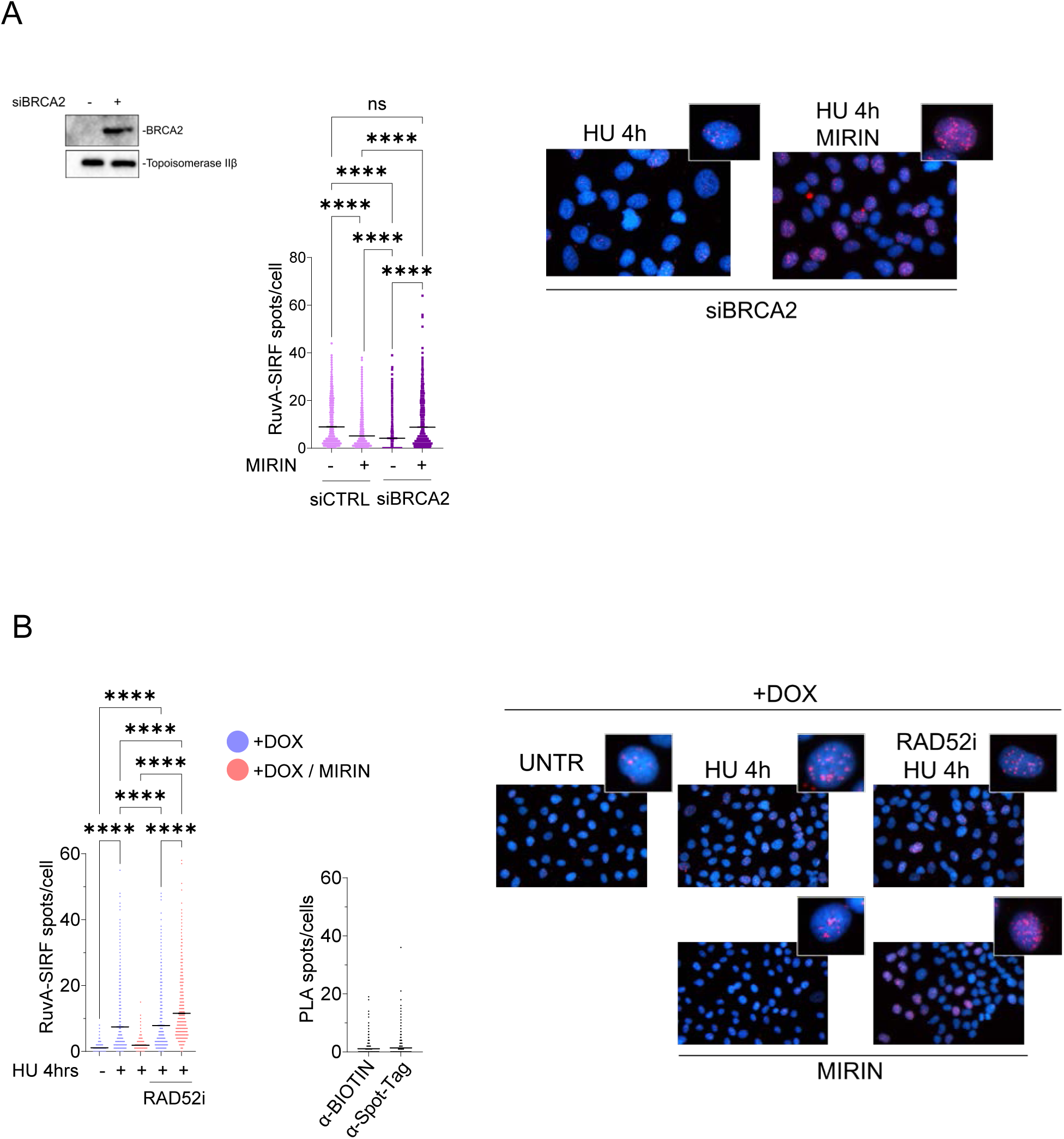
Accumulation of RuvA at perturbed forks requires fork reversal mechanisms. **(A)** Analysis of RuvA-Spot recruitment at nascent DNA by SIRF in cells transfected with siRNA targeting *BRCA2*. Western blot (top) confirms protein knockdown; Topoisomerase IIβ served as a loading control. Cells were treated as indicated; where specified, the MRE11 inhibitor MIRIN was added 30 min prior to treatment. Graph shows quantification of PLA spots per cell (> 1000 cells per condition; N = 3). Data includes mean ± SEM; \*\*\*\**PP* < 0.0001 by Kruskal–Wallis test. Representative images are shown. **(B)** Evaluation of RuvA recruitment at perturbed forks following inhibition of RAD52 and MRE11. Cells were treated as indicated; RAD52 inhibitor (RAD52i/EGC) and MIRIN were added 30 min prior to RS induction. Graph shows the number of PLA spots per cell and the mens ± SEM (> 1000 cells per condition); \*\*\*\**PP* < 0.0001 by Kruskal–Wallis test. Negative controls (single-antibody incubations with anti-biotin or anti-Spot-Tag) and representative images are included.

Our previous findings show that RAD52 structurally competes with SMARCAL1 at perturbed replication forks and that RAD52 inhibition or depletion induces MRE11-dependent fork degradation presumably because of excessive SMARCAL1 localisation at fork and increased frequency of fork reversal following HU exposure ^23,24^. Thus, we used our RuvA biosensor to evaluate the dynamics of RFs under these conditions. Quantitative analysis of the SIRF assay in the U2OS inducible model treated with the RAD52 inhibitor EGC ^48^ demonstrated that RuvA recruitment was enhanced in this context compared to exposure to HU alone (Fig. 3B), suggesting that the number of RFs in RAD52-inhibited cells is increased as compared with the wild-type. However, when cells were treated with MIRIN, the RuvA-Spot SIRF signals was further significantly increased over the wild-type (Fig. 3B), supporting the hypothesised mechanism and a deregulated formation of four-way junctions, likely RFs.

Taken together, these data indicate that RuvA specifically binds to reversed forks and that experimental variations in the frequency of fork reversal directly modulate the recruitment of RuvA either in down or in up. Moreover, using RuvA as a RF biosensor, we confirmed that inhibition of RAD52 stimulates formation of these structures at stalled forks followed by rapid degradation by MRE11.

### RuvA detects RAD51-dependent four-way junction accumulation in nucleoli after fork stalling

Following our demonstration that RuvA localizes at reversed forks by SIRF assay, we sought to determine if it could be used also by immunofluorescence (IF) confocal microscopy. Our initial IF analysis showed that RuvA cannot be detected directly using GFP. Thus, we used the RuvA-Spot inducible cells and the anti-Spot-Tag antibody for detection. As shown in Figure 4A, IF failed to evidence clear RuvA foci after 4 hrs of 2 mM HU but revealed a distinct staining pattern that was similar, in morphology, to that of the nucleoli. Of note, this pattern was detected also in untreated cells but with a less intense staining (Fig. 4A). To investigate this further, we performed a co-immunofluorescence assay in the RuvA-Spot inducible cells and used Nucleolin as a canonical nucleolar marker. As expected, nucleolar-like localisation of RuvA-Spot was readily detected in untreated cells although the number of positive cells and the intensity of the immunofluorescence increased in response to HU or CPT (Fig. 4C; histogram and microscopy fields). Since our data indicate that localisation of RuvA at forks is dependent on RAD51 and RAD51 is key in the formation of four-way junctions, we combined RS with RAD51i to analyse the effect on the nucleolar-like localisation. Inhibition of RAD51 reduced the number of RuvA-Spot-positive cells and, most interestingly, suppressed the RS-induced increase in the intensity of the signal (Fig. 4C and D). As shown by Figure 4D and E, RuvA specifically co-localizes completely with Nucleolin, establishing its presence within this nucleolar compartment while no difference was observed in the percentage of Nucleolin-positive cells irrespective of the expression of RuvA-Spot (Fig. 4F). Furthermore, we observed that this localization is dynamically maintained following cellular exposure to RS induced by HU or a low-dose of CPT (Fig. 4C-E). Notably, the inhibition of RAD51 significantly disrupted this nucleolar co-localization, suggesting a regulated recruitment or retention mechanism (Fig. 4C-E; histograms and microscopy fields).

**Figure 4.**
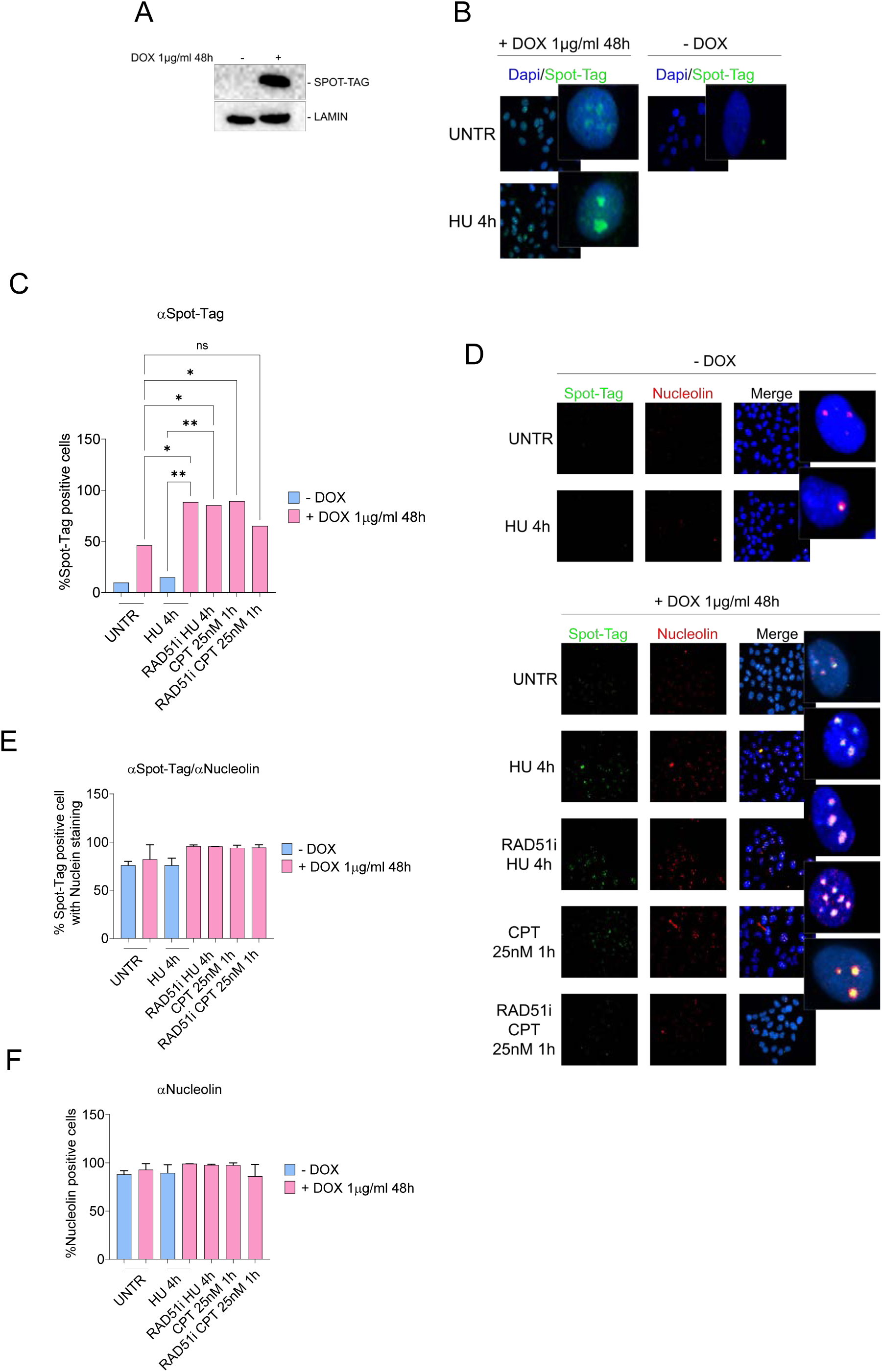
RuvA accumulates in nucleolar structures under replication stress. **(A)** Western blot analysis of RuvA-Spot-Tag expression control (± 1 *μμ*g/ml DOX for 48 hrs) with Lamin as a loading control. **(B)** Immunofluorescence (IF) analysis of RuvA-Spot-Tag reveals focal localization in large punctate nuclear structures consistent with nucleoli. Treatment with 2 mM hydroxyurea (HU) for 4 h increases puncta intensity, indicating enhanced recruitment under replication stress. **(C-F)** Cells were left untreated or treated with 2 mM HU (4 hrs) or 25 nM camptothecin (CPT, 1 hr), with or without a 30 min pre-treatment with 27 *μμ*M RAD51 inhibitor (B02). In the graphs are shown: percentage of RuvA-Spot-Tag-positive cells (C); percentage of RuvA-Spot-Tag-positive cells colocalizing with Nucleolin (E); percentage of Nucleolin-positive cells (F). Data represent mean ± SD from three independent experiments (at least 100 cells per replicate). Statistical significance was calculated using one-way ANOVA (*PP* < 0.01, \**PP* < 0.05, ns = not significant). **(D)** Representative images from co-immunofluorescence (Co-IF) analysis of RuvA-Spot-Tag (green) and Nucleolin (red).

Taken together, our data demonstrate that RuvA recognizes four-way junction accumulating in the nucleoli and that these structures can increase upon RS in a RAD51-dependent manner.

### Ectopic RuvA binds firmly four-way junctions and can be used to investigate consequences of their long-term persistence

In bacteria, RuvA is an avid four-way junction binder and is displaced after RuvC-dependent cleavage ^12,41^. When expressed in cells, RuvA only binds these DNA structures acting as a biosensor by exploiting its strong affinity. Thus, we speculated that our approach could be used not just for visualisation but also for mechanistic investigation of the consequences of long-term freezing of four-way junctions, especially of RFs under perturbed replication. To this end, we assessed if RuvA-Spot could persist at RFs or at four-way junctions at large, during recovery from RS. To this end, we analysed the presence of RuvA by SIRF during recovery from the perturbation of DNA replication induced by the low dose of CPT, which is a stronger inducer of RFs. As shown in Figure 5A, RuvA was readily detected at perturbed replication forks by SIRF but did not significantly reduce binding during recovery. In contrast, localisation of SMARCAL1 at perturbed forks greatly decreased with recovery (Fig. 5A). Subsequently, we investigated if its expression affects cellular replication and whether it triggers an increase in DNA damage. To address this, we employed DNA fibre assays to measure fork progression speed at a single-molecule level. Cells were induced to express RuvA-Spot and 48 h thereafter their active forks were labelled with CldU. After washing, fork arrest was induced by 2 mM HU followed by a recovery in IdU to label restarting forks. As expected, cells efficiently recovered replication from stalled forks (Fig. 5B). In contrast, cells expressing RuvA restarted less efficiently replication after recovery from HU (Fig. 5B).

**Figure 5.**
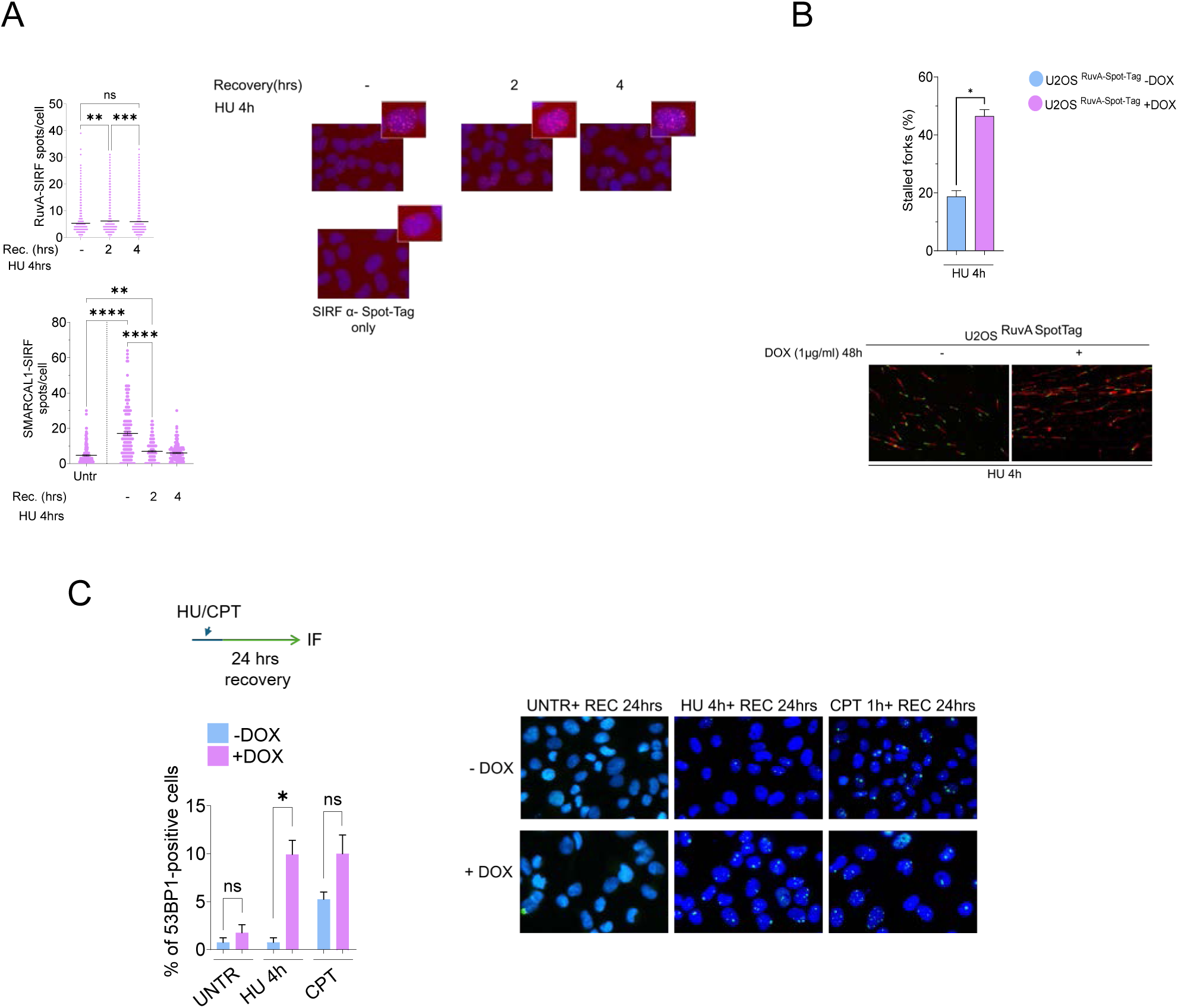
RuvA dynamics delay replication fork restart by binding reversed forks. **(A)** Kinetics of RuvA-Spot and SMARCAL1 recruitment following replication stress. Cells were treated with 2 mM HU for 4 hrs and allowed to recover in fresh DMEM for the indicated time points. Graph shows PLA spots per cell (> 1000 cells per condition; N = 3). Single-antibody incubations (anti-Spot-Tag only) served as negative control for RuvA. Data represent individual values and indicate mean ± SEM; ns = not significant, \*\**P* < 0.01, \*\*\**P* < 0.001 by Kruskal–Wallis test. Representative images are shown. **(B)** Analysis of replication fork restart by DNA fiber assay. Experimental labeling scheme is depicted above. Graph shows the percentage of stalled replication forks from N = 3. Representative DNA fiber tracks are shown. \*\*\**P* < 0.001 by one-way ANOVA. **(C)** Analysis of 53BP1 foci formation following recovery. Cells were treated as indicated, released into fresh DMEM for 24 h, extensively washed with PBS, and immunostained for 53BP1. Graph shows the percentage of 53BP1 NB-positive cells. Data represent mean ± SEM from N = 4; ns = not significant, \**P* < 0.05 by Mann-Whitney test.

We subsequently analysed the accumulation of DNA damage following RuvA expression in comparison to wild-type cells to have preliminary proof of the effects of “freezed” RFs. This was evaluated through the accumulation of 53BP1 NBs as a recognized readout of cellular stress and damage. To this end, cells were induced to express RuvA-Spot with Dox for 48 hrs and treated with 2 mM HU 4 hrs or 25 nM CPT 1h before being recovered in drug-free medium for 24 hrs. At the end of the recovery, cells were immunostained with the anti-53BP1 antibody and counted for the presence of NBs. As show in Figure 5C, cells expressing RuvA-Spot showed an increased percentage of 53BP1-positive cells as compared with control cells (-Dox). In cells recovering from HU, expression of RuvA-Spot resulted in about 4-fold more 53BP1-positive cells while the difference was less noticeable after recovery from 25 nM CPT, although in this case the intensity of the staining conferred in positive cells appeared increased. Despite in CPT-treated cells the differences were not statistically significant because of the few cells showing 53BP1 NBs they were reproducible among replicates.

Altogether, our findings demonstrate that ectopic RuvA-Spot persists at four-way junctions in the cells and suggest that, in human cells, persistence of four-way junctions – likely RFs – at perturbed replication forks affects replication fork recovery and results in a small but reproducible increase in the frequency of 53BP1 NBs.

## DISCUSSION

In this study, we investigated the potential of using the bacterial protein RuvA to detect the presence of RFs and four-way junctions at large in human cells. RuvA is a component of the bacterial resolvase RuvABC that recognizes and binds four-way structures like HJs. Our previous work demonstrated that bacterial RuvA can be expressed in human cells and can bind DNA – likely HJs – but only using biochemical assay ^44^ and without addressing the identity of the structure or the suitability for imaging at the ingle-cell level.

Here, using two independent variants of RuvA, specifically assembled for expression in the nucleus and ease of labelling with validated antibodies, we report that RuvA can be used, when expressed in human cells, as an imaging biosensor to determine the presence of RFs or, more generally, four-way junctions in vivo. Moreover, we provide proof of principle that ectopic expression of RuvA can be also used as a tool to investigate the response of the cells to persisting four-way junctions. A similar approach has been successfully exploited for detection of R-loops by a catalytic-dead RNaseH1 ^40^, however, R-loops can be visualized with all the required controls, also by the S9.6 antibody ^49^ while our tool represents the first validated approach to image RFs and HJs *in vivo*. In the past, other bacterial proteins have been used to handle HJs in human cells. For instance, ectopic RuvC or RusA have been used to eliminate four-way junctions addressing the correlation between a pathological phenotype and their accumulation ^50–52^. Our approach inverts this logic and using the binding module of RuvABC alone developed a true biosensor with the additional ability to act as a stabilizer of these structures.

Interestingly, while we failed to detect RuvA foci under RS we readily observed binding at nascent DNA of perturbed forks by SIRF. These results are not at odds and could imply that RFs do not form clusters in the nucleus, as replication factories do, and that single tetramers of RuvA recognise the structure, as expected. The presence of isolated RFs or HJs after fork stalling would likely prevent detection either by GFP or by antibody, requiring an approach like SIRF based on signal amplification. Our observations can likely suggest that any other engineered bacterial HJ-binder, such as catalytic-dead RuvC or RusA, could be used as biosensor if coupled with SIRF.

The use of SIRF to detect RuvA however comes with additional advantages, such as the specific visualization at RFs or HJs at/near the stalled fork. Interestingly, the results obtained with our RuvA biosensor recapitulate well what reported using EM. Indeed, the kinetics of accumulation of RuvA fits well that reported by EM in CPT or HU-treated cells ^16,26^. Moreover, the dependency of RuvA localisation at perturbed forks from SMARCAL1, RAD51 or ZRANB3 and the extent of suppression derived from single or combined inactivation mirrors what reported previously by EM, with SMARCAL1 and RAD51 being key for RuvA localisation ^29,31^. The consistent observation with EM, the shared genetic dependencies and effect of the HJ-binding mutant all support RuvA as a valid biosensor of RFs and four-way junctions *in vivo*.

Not only our data were supported by evidence that the downregulation of fork reversal through SMARCAL1 depletion or RAD51 inhibition reduced RuvA recruitment to DNA. Conversely, visualisation of RuvA by SIRF also recapitulated the established reduction of RFs in BRCA2-depleted cells ^29,31,34^, also correctly reporting their restoration upon inhibition of MRE11 ^29^. Moreover, using RuvA-SIRF as a RF biosensor, we have been able to detect the increasing fork reversal via RAD52 inhibition speculated in our previous work ^24^.

Although standard immunofluorescence analysis failed to detect RuvA foci in treated cells, it revealed that RuvA localizes in nucleoli, which was confirmed by co-immunofluorescence with nucleolin. This is striking for two reasons: first because any RuvA-SIRF cluster was apparent to accumulate at nucleoli; SIRF relies on the presence of nascent DNA reporting interactions at the fork thus nucleolar staining of RuvA is possibly outside replication forks. Second because localisation fades when RAD51 is inhibited preventing formation of four-way junctions – likely HJs. Nucleoli are well-known as site where recombination is elevated ^53^ and our results are consistent with this feature.

In addition to be a biosensor for RFs and four-way junctions in general, RuvA is also a tool to pinpoint the cellular consequences to persistence of these structures after fork perturbations. Indeed, the strong avidity of RuvA for four-way junctions leads to persistence at replicating sites after withdrawal of the RS. Using RuvA-Spot for this purpose, our data give proofs for a pathological consequence of persistent RFs/four-way junctions in the cell. Interestingly, the reduced recovery of stalled forks in the presence of RFs is consistent with the expected requirement of their resolution for a timely resumption of replication ^27^. This supports the need - in mechanistic studies - of a tuneable RuvA expression system like that we developed in U2OS cells. Indeed, sustained overexpression after transient transfection can stimulate micronuclei in specific genetic backgrounds ^44^.

Overall, our findings demonstrate that RuvA can be utilized as a sensor for reversed forks and four-way junctions in general, which is a key development for studying cellular genome instability. The ability to investigate whether genomic instability is linked to the abundance of reversed forks makes RuvA a valuable tool for studying mechanisms frequently associated with diseases and tumours. Moreover, further improvements of our tool could allow its exploitation for the characterisation of hot-spots of RFs accumulation, landscape of associated factors and identification of genetic dependencies using ad-hoc screens.

## ACKNOWLEDGMENTS

The authors want to thank Prof. Massimo Lopes for discussion.

AUTHOR CONTRIBUTIONS: E.M. performed all the cell biology experiments for functional characterisation of the biosensor. E.R. performed the characterization of nucleolar localisation and contributed to biochemical experiments. E.M., E.R. analysed data and contributed to designing the experiments and writing the paper. P.P. contributed to analyse data. A.F., and P.P designed experiments, supervised research and wrote the paper. All authors approved the paper.

