## Supplementary Figures and Legends for "A novel imaging biosensor for the detection of reversed replication forks and four-way junctions in human cells"

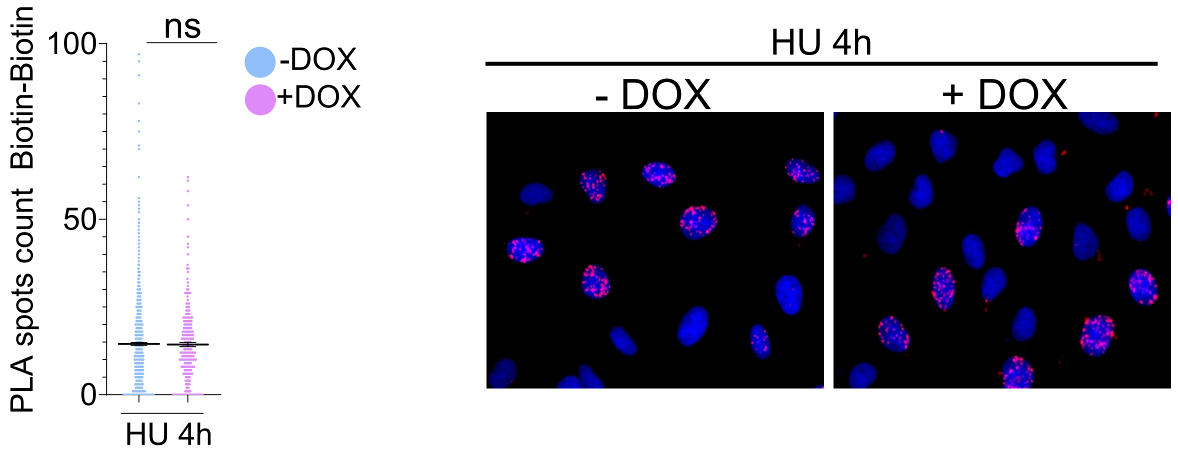


**Supplementary Figure 1**. After 48 hrs of doxycline induction cells were labelled with EdU to detect S-phase and treated as indicated. The graph shows the number of PLA spot per nucleus in S-phase. The images were acquired and analysed by Olympus ScanR High Content Imaging System from 3 combined experiments. All the values above include means ± SEM (ns = not significant; ****P < 0.001; Kruskal–Wallis test). Representative images are shown.

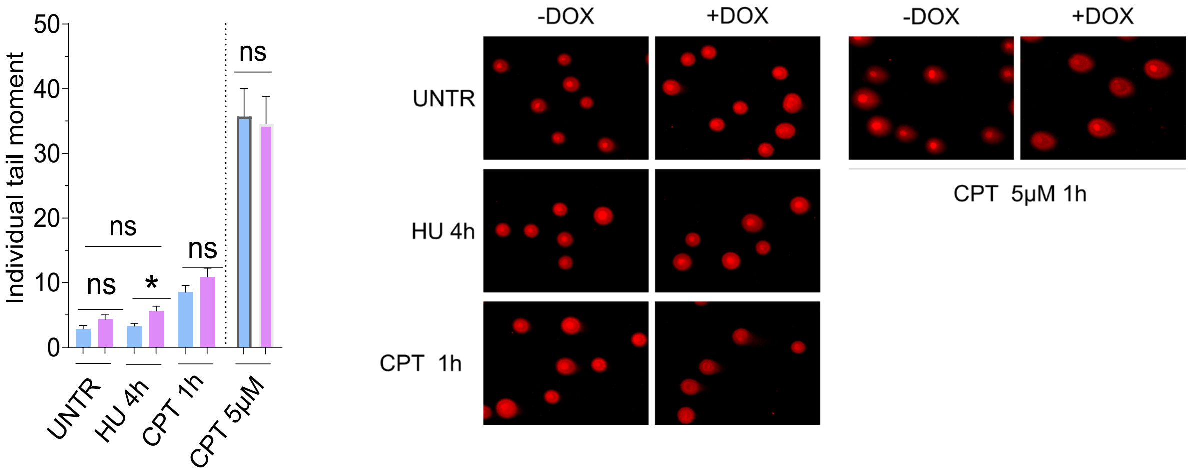


**Supplementary Figure 2**. After 48 hrs of doxycycline induction cells were treated as indicated. Cellular pellets were collected and used to perform neutral Comet assay. Representative Images are shown. Data are presented as individual tail moment values from N=3 with at least 90 comets/each. The mean of tail moment ±SEM is indicated (ns = not significant; * P < 0.5; Mann–Whitney test)
